# Potential Autoimmune Association between Benign Paroxysmal Positional Vertigo and Immune-mediated Skin Conditions: Population-based Cohort Study

**DOI:** 10.1101/676973

**Authors:** Hyung Jin Hahn, Sang Gyu Kwak, Dong-Kyu Kim, Jong-Yeup Kim

**Affiliations:** Department of Dermatology, College of Medicine, *Konyang* University, *Daejeon*, Republic of Korea; School of Medicine, Department of Medical Statistics, Catholic University of *Daegu*; Department of Otorhinolaryngology-Head and Neck Surgery, *Chuncheon* Sacred Heart Hospital; Institute of New Frontier Research, *Hallym* University College of Medicine, *Chuncheon*, Republic of Korea; Department of Otorhinolaryngology–Head and Neck Surgery, College of Medicine, *Konyang* University, *Daejeon*, Republic of Korea; Department of Biomedical Informatics, College of Medicine, *Konyang* University, *Daejeon*, Republic of Korea; *Myunggok* Medical Research Institute, College of Medicine, *Konyang* University, *Daejeon*, Republic of Korea

**Author notes:** **Corresponding authors:** Dong-Kyu Kim, MD, Ph.D, Department of Otorhinolaryngology-Head and Neck Surgery, *Chuncheon* Sacred Heart Hospital, *Hallym* University College of Medicine (200-704), 77, *Sakju-ro, Chuncheon-si, Gangwon-do*, Republic of Korea, Jong-Yeup Kim MD, Ph.D, Department of Biomedical Informatics, College of Medicine, *Konyang* University, 158 *Gwanjeodong-ro, Seo-gu, Daejeon* 35365, Republic of Korea.

**Keywords:** Benign Paroxysmal Positional Vertigo (BPPV), atopic dermatitis, vitiligo, autoimmunity, big data

## Abstract

**Background:** Benign paroxysmal positional vertigo (BPPV), an idiopathic disorder of sudden sensorineural hearing loss and vertigo, shares many similarities with two common skin conditions, atopic dermatitis (AD) and vitiligo. Recent studies have suggested that BPPV may be related to or triggered by autoimmune conditions, notably hypothyroidism and giant cell arteritis (GCA).

**Objective:** These evidences prompted the authors to entertain the possibility of immunological bridge between BPPV and the two skin conditions. The authors have tested this hypothesis with population-based cohort from the National Health Insurance Service Database of Korea.

**Methods:** A cohort of 1.1 million patients was extracted from the DB. Using *χ*^2^ tests, prevalence of the two skin disorders in terms of BPPV status was analysed.

**Results:** In AD patients, the prevalence of BPPV was 30% lower, while there was no statistically significant relationship between BPPV prevalence and vitiligo. The relationship between vitiligo and BPPV was significant in younger subgroup only. Socio-economic subgroup analysis revealed the observed patterns are primarily a middle-upper class phenomenon.

**Limitations:** Uncertainty regarding temporal sequence of onset, and lack of detail on disease severity and subtype might have kept the authors from drawing more refined conclusion.

**Conclusion:** AD and vitiligo might be linked to BPPV through the action of certain components of cellular immunity, but follow-up studies based on large population cohort would be needed to add more substance to our findings.

## Introduction

Benign paroxysmal positional vertigo (BPPV), along with *Ménière* disease (MD) and labyrhinthitis, is one of the most commonly encountered forms of vertiginous disorder. Accounting for almost 50% of all individuals with peripheral vestibular dysfunction [1], one-year and lifetime prevalence of this largely idiopathic entity is believed to be around 1.6% and 2.4% respectively, with a slight predilection for females [2,3,4]. The vertigo of sudden onset, that is characteristic of BPPV, is thought to be instigated by displacement of inner ear crystals (otoliths), with the stones lodging within the lumen of the semi-circular canals [5], which in turn culminates in perturbed rotational balance. While postural or head position changes as precipitating factors of BPPV symptoms is reasonably well established, whether there are other factors accounting for the stimulation of labyrinthine receptors is a matter of ongoing debate. Traditional views have held that the phenomenon is primarily of mechanical nature, while some schools of thought have embraced the notion of autoimmune etiopathogenesis for BPPV. Studies over the past several years have reinforced the notion of the endolymphatic sac, where these “ear rocks” originate, as a vibrant hub of immunological interactions, rather than an inert structural locale [6]. The labyrinthine sac is known to harbour inner ear autoantigens of various molecular weights, which can trigger type II (surface antigens) or type III (antigen-antibody immunocomplexes) hypersensitivities [7,8]. One facet of the autoimmune hypothesis points to this type of immune-complex formation and diffusion of these complexes into the inner ear surface, as a direct cause of the triggering of the symptoms [9]. For the past couple of decades, there has been some debate raging over the issue of thyroid status with respect to BPPV pathogenesis. Some authors have found that there is a positive correlation between the chronic inner ear disorder and Hashimoto thyroiditis, which is the most common cause of hypothyroidism [10,11]. Furthermore, Papi *et al*. showed that BPPV is related to autoimmune chronic thyroiditis (ACT), and that the association is independent of thyroid status [12]. However, despite these reports, the issue remains unsettled. To explore the issue from a different perspective, the authors have tested how the chronic inner ear condition is related to two of the most prevalent inflammatory skin diseases, atopic dermatitis and vitiligo. The strong ties of these two conditions to a variety of autoimmune diseases are fairly well established, and in particular, their relationship to thyroid diseases has been explored in a wealth of previous literatures [13,14]. Also, because each of the two entities has distinctive immuno-pathological mechanism, they were deemed suitable for the purposes of comparison with BPPV.

The aim of the authors in the present investigation was to offer an alternative and complementary explanation for the controversial role of autoimmunity in BPPV, through analysis based on population-based cohorts from the National Health Insurance Service of Korea (NHIS) database.

## Results

### Baseline characteristics

Baseline demographic information is summarized in Table 1. The entire study cohort was made up of 1,113,656 individuals, with nearly equal sex distribution (M:F=50.1:49.9). Geographically, the highest proportion of the cohort was drawn from Seoul and *Gyeonggi* Province, a metropolitan area surrounding the nation’s capital (at around 21% apiece). Other major cities and the metropolitan provinces of the country were evenly represented. Each eligible citizen is subject to either one of the two coverage plans, *i.e*., employee-insured or self employee-insured, or when applicable, the Korean Medical Aid program. The cohort was divided into ten income brackets (deciles), and then regrouped as *lower* (brackets 1 through 4), *middle* (brackets 5 through 7), or *upper* (brackets 8 through 10) income tiers. The study cohort was also divided from grade of 0 to 6 according to the extent of their disability. For the entire cohort, the “baseline” prevalence of BPPV, AD, and vitiligo were computed at 0.69%, 0.72% and 0.11%, respectively.

**TABLE 1.** Baseline characteristics. *Abbreviations*: *AD*, atopic dermatitis; *BPPV*, benign paroxysmal positional vertigo.

| Variable |  | *n* | % |
| --- | --- | --- | --- |
| Total patients |  | 1,113,656 | 100.0 |
| Sex | Male | 558,186 | 50.12 |
|  | Female | 555,470 | 49.88 |
| Age distribution (years) | 0 | 108,614 | 9.75 |
|  | 1~4 | 52,082 | 4.68 |
|  | 5~9 | 74,637 | 6.70 |
|  | 10~14 | 70,962 | 6.37 |
|  | 15~19 | 70,135 | 6.30 |
|  | 20~24 | 84,128 | 7.55 |
|  | 25~29 | 85,303 | 7.66 |
|  | 30~34 | 97,177 | 8.73 |
|  | 35~39 | 88,505 | 7.95 |
|  | 40~44 | 93,776 | 8.42 |
|  | 45~49 | 72,811 | 6.54 |
|  | 50~54 | 53,333 | 4.79 |
|  | 55~59 | 43,015 | 3.86 |
|  | 60~64 | 42,753 | 3.84 |
|  | 65~69 | 31,891 | 2.86 |
|  | 70~74 | 20,939 | 1.88 |
|  | 75~79 | 12,850 | 1.15 |
|  | 80~84 | 7,063 | 0.63 |
|  | 85+ | 3,682 | 0.33 |
| Age group (years) | ≤65 | 1,037,231 | 93.14 |
|  | >65 | 76,425 | 6.86 |
| Income deciles | 0 | 28,332 | 2.54 |
|  | 1 | 63,625 | 5.71 |
|  | 2 | 65,585 | 5.89 |
|  | 3 | 77,713 | 6.98 |
|  | 4 | 91,386 | 8.21 |
|  | 5 | 105,376 | 9.46 |
|  | 6 | 119,392 | 10.72 |
|  | 7 | 131,489 | 11.81 |
|  | 8 | 142,484 | 12.79 |
|  | 9 | 145,778 | 13.09 |
|  | 10 | 142496 | 12.80 |
| Income Ranges | 0~4 | 326,641 | 29.33 |
|  | 5~7 | 356,257 | 31.99 |
|  | 8~10 | 430,758 | 38.68 |
| Disability Grade | Normal (Grade 0) | 1,087,242 | 97.63 |
|  | Moderate (Grade 1~2) | 8,943 | 0.80 |
|  | Severe (Grade 3~6) | 17,471 | 1.57 |
| BPPV | No | 1,105,918 | 99.31 |
|  | Yes | 7,738 | 0.69 |
| AD | No | 1,105,616 | 99.28 |
|  | Yes | 8,040 | 0.72 |
| Vitiligo | No | 1,112,385 | 99.89 |
|  | Yes | 1,271 | 0.11 |

**TABLE 2.** Relationship between BPPV and atopic and vitiligo. *Abbreviations*: *AD*, atopic dermatitis; *BPPV*, benign paroxysmal positional vertigo. *statistically significant for *p*<0.05.

| Variable |  | BPPV, n (%) |  | p-value |
| --- | --- | --- | --- | --- |
|  |  | No | Yes |  |
| AD | No | 1,097,917 (99.28) | 7,699 (99.50) | 0.023* |
|  | Yes | 8,001 (0.72) | 39 (0.50) |  |
| Vitiligo | No | 1,104,658 (99.89) | 7,727 (99.86) | 0.464 |
|  | Yes | 1,260 (0.11) | 11 (0.14) |  |

**TABLE 3.** Relationship between BPPV and AD and vitiligo by sex. *Abbreviations*: *AD*, atopic dermatitis; *BPPV*, benign paroxysmal positional vertigo. *statistically significant for *p*<0.05.

| Sex | Variable |  | BPPV, *n* (%) |  | *p*-value |
| --- | --- | --- | --- | --- | --- |
|  |  |  | No | Yes |  |
| Male | AD | No | 552,247 (99.34) | 2,284 (99.61) | 0.119 |
|  |  | Yes | 3,646 (0.66) | 9 (0.39) |  |
|  | vitiligo | No | 555,361 (99.90) | 2,291 (99.91) | 0.896 |
|  |  | Yes | 532(0.10) | 2(0.09) |  |
| Female | AD | No | 545,670 (99.21) | 5,415 (99.45) | 0.045* |
|  |  | Yes | 4,355 (0.79) | 30 (0.55) |  |
|  | Vitiligo | No | 549,297 (99.87) | 5,436 (99.83) | 0.507 |
|  |  | Yes | 728 (0.13) | 9 (0.17) |  |

**TABLE 4.** Relationship between BPPV and atopic and vitiligo by age (≤65 & >65). *Abbreviations*: *AD*, atopic dermatitis; BPP*V*, benign paroxysmal positional vertigo. *statistically significant for *p*<0.05.

| Age group | Variable |  | BPPV, *n* (%) |  | *p*-value |
| --- | --- | --- | --- | --- | --- |
|  |  |  | No | Yes |  |
| ≤65 | AD | No | 1,022,463 (99.23) | 6,848 (99.49) | 0.015* |
|  |  | Yes | 7,885 (0.77) | 35 (0.51) |  |
|  | Vitiligo | No | 1,029,167 (99.89) | 6,873 (99.85) | 0.454 |
|  |  | Yes | 1,181 (0.11) | 10 (0.15) |  |
| >65 | AD | No | 75,454 (99.85) | 851 (99.53) | 0.021* |
|  |  | Yes | 116 (0.15) | 4 (0.47) |  |
|  | Vitiligo | No | 75,491 (99.90) | 854 (99.88) | 0.911 |
|  |  | Yes | 79 (0.10) | 1 (0.12) |  |

**TABLE 5.** Relationship between BPPV and atopic and vitiligo by income tiers. *Abbreviations*: *AD*, atopic dermatitis; *BPPV*, benign paroxysmal positional vertigo. *statistically significant for *p*<0.05.

| Income Brackets | Variable |  | BPPV, *n* (%) |  | *p*-value |
| --- | --- | --- | --- | --- | --- |
|  |  |  | No | Yes |  |
| 0~4<br>[Lower] | AD | No | 322,809 (99.48) | 2,116 (99.39) | 0.585 |
|  |  | Yes | 1,703 (0.52) | 13 (0.61) |  |
|  | Vitiligo | No | 324,223 (99.91) | 2,123 (99.72) | 0.003* |
|  |  | Yes | 289 (0.09) | 6 (0.28) |  |
| 5~7<br>[Middle] | AD | No | 351,513 (99.26) | 2,130 (99.58) | 0.089 |
|  |  | Yes | 2,605 (0.74) | 9 (0.42) |  |
|  | Vitiligo | No | 353,754 (99.90) | 2,138 (99.95) | 0.419 |
|  |  | Yes | 364 (0.10) | 1 (0.05) |  |
| 8~10<br>[Upper] | AD | No | 423,595 (99.14) | 3,453 (99.51) | 0.017* |
|  |  | Yes | 3,693 (0.86) | 17 (0.49) |  |
|  | Vitiligo | No | 426,681 (99.86) | 3,466 (99.88) | 0.676 |
|  |  | Yes | 607 (0.14) | 4 (0.12) |  |

**TABLE 6.** Relationship between BPPV and atopic dermatitis/vitiligo by disability. *Abbreviations*: *AD*, atopic dermatitis; *BPPV*, benign paroxysmal positional vertigo. *statistically significant for *p*<0.05.

| Grade | Variable |  | BPPV, *n* (%) |  | *p*-value |
| --- | --- | --- | --- | --- | --- |
|  |  |  | No | Yes |  |
| Normal | AD | No | 1,071,769 (99.27) | 7,503 (99.48) | 0.027* |
|  |  | Yes | 7,931 (0.73) | 39 (0.52) |  |
|  | Vitiligo | No | 1,078,459 (99.89) | 7,531 (99.85) | 0.430 |
|  |  | Yes | 1,241 (0.11) | 11 (0.15) |  |
| Moderate | AD | No | 8,876 (99.74) | 44 (100.0) | 0.736 |
|  |  | Yes | 23 (0.26) | 0 (0.0) |  |
|  | Vitiligo | No | 8,897 (99.98) | 44 (100.0) | 0.921 |
|  |  | Yes | 2 (0.02) | 0 (0.0) |  |
| Severe | AD | No | 17,272 (99.73) | 152 (100.0) | 0.521 |
|  |  | Yes | 47 (0.27) | 0 (0.0) |  |
|  | Vitiligo | No | 17,302 (99.90) | 152 (100.0) | 0.699 |
|  |  | Yes | 17 (0.10) | 0 (0.0) |  |

### Prevalence of BPPV in terms of skin disease status

The prevalence of BPPV in AD individuals was 0.72%. In non-AD patients, the prevalence was 0.50%. The finding was statistically significant (*p*=0.023). The prevalence of BPPV in vitiligo and non-vitiligo individuals was 0.11% and 0.14%, respectively. However, these relationships were not statistically significant (*p*=0.464).

### Sex and age

A similar pattern prevailed after male and female cohorts were considered separately, although the relationship between BPPV and AD was not statistically significant in males (*p*=0.119). Age group analysis revealed that the higher prevalence of vitiligo in BPPV patients held true only for the younger individuals (*p*=0.000). Other findings were not statistically significant.

### Socioeconomic subgroups

On χ^2^ test, the BPPV-vitiligo relationship was valid only in the “middle” income tier (brackets 5 through 7, *p*=0.000). On the other hand, the lower BPPV prevalence in AD patients was seen only in the “upper” tier (brackets 8 through 10, *p*=0.037). Other relationships were not statistically significant.

### Disability

The BPPV-AD/vitiligo relationship was also analysed by disability status. χ^2^ analysis revealed virtually the same results for individuals without disability as the whole cohort (*p*-values of 0.026 and 0.000 for AD and vitiligo, respectively). Interestingly, there was a reversal of pattern in the subgroup with moderate (grade 1 & 2) disability, with a five-fold increase in AD prevalence in BPPV patients (1 of 56 *versus* 22 of 8,864 in non-BPPV patients; *p*=0.025).

## Discussion

While many causes are cited as possible players in its aetiology, nearly 60% of BPPV cases are still idiopathic [15], and it would not be completely implausible that at least a part of this proportion may be attributed to autoimmune causes. The autoimmune aspect of BPPV pathogenesis is a relatively recent development, and it is not “officially” recognised and given serious consideration in most established text sources. Apart from the issue involving thyroid, another piece of evidence in favour of the autoimmunity theory may be the link between BPPV and giant cell arteritis (GCA) [16]. The pathophysiology of GCA involves dendritic cells (DC) in the vessel wall which attract T cells and macrophages to form granulomatous infiltrates. T helper 17 cells (Th 17), interconnected with interleukin (IL) 6, IL-17 and IL-21, play an essential role, and this pathway can be blocked by corticosteroids [17]. From the population-based cohort of the present study, it was revealed that individuals with AD, or history of AD (BPPV tends to affect senile individuals whereas the incidence of AD peaks at late-20’s to mid-30’s) is about 30% less likely to be affected with BPPV. Atopic dermatitis is a chronic, inflammatory disorder of the skin, in which thickening of the epidermis is one of the hallmarks [18]. The epidermal hyperplasia results from complex network of interactions between keratinocytes and T-cells, mediated by several cytokines and chemokines [19]. Several studies from recent years have found that systemic inflammatory disorders are more prone to developing inner ear diseases; in psoriasis, which is also a chronic inflammatory disease characterized by hyperplastic epidermis, there is an increased risk of sudden sensorineural hearing loss [20,21]. Given the systemic nature of AD inflammation [22], it is likely that the surface of the labyrinthine sac of the utricle is also affected in a way that secures the position of otoliths within the utricle. Meanwhile in subgroup analysis. there was a sharp contrast between younger and senile cohorts. The higher prevalence of BPPV in senile patients with AD may suggest that immune senescence plays a role in the pathogenesis of BPPV [23]. In the sub-analysis by socioeconomic status, the prevalence of vitiligo was three times higher in BPPV individuals that belong to the lower income tier (deciles 1 through 4). On the other hand, for the upper tier subgroup (deciles 8 through 10), the prevalence of AD in BPPV individuals was nearly one-half that of the non-BPPV cohort. This phenomenon may reflect unequal distribution of healthcare utilization across the socioeconomic hierarchy. Individuals lacking financial means and leisure are less likely to seek medical care for cutaneous signs of vitiligo at secondary or higher referral centres. However, they would be more compelled to take the trouble for symptoms of BPPV, which can seriously undermine the individual’s day-to-day operation, and at larger centres this in turn would likely lead to the diagnosis of concurrent vitiligo.

The authors acknowledge that this study was subject to some limitations. Due to its cross-sectional nature, temporal relations between BPPV and skin disorders could not be established. Also, lack of information regarding disease severity and subtype (*e.g.*, anterior or horizontal canal *versus* posterior canal BPPV [24,25]) had impeded more detailed analysis, which would have allowed the authors to propose more elaborate disease mechanism. Finally, it was revealed by subgroup analysis that the relationship between the skin conditions and BPPV was statistically significant only for individuals *without* disability.

In conclusion, the present study represents a unique attempt to form a potential link between autoimmune conditions and BPPV, which has been considered to originate from mechanical/physical causes for the most part. Despite a few shortcomings, the study allowed the authors to glimpse through the underlying patho-mechanism of three puzzling conditions, and on the process gain some unique insights and perspectives. Further studies, with more thorough and sophisticated databases, would enable us to build upon this ground and yield more refined conclusions, including its therapeutic implications.

## Methods

### Database (DB)

All study conduct adhered to the tenets of the Declaration of Helsinki. This study utilized KNHIS-NSC data (NHIS-2018-2-252), made by the National Health Insurance Service (NHIS) and was approved by the Institutional Review Board (IRB) of *Hallym* Medical University *Chuncheon* Sacred Hospital (IRB No. 2018-08-018). The need for written informed consent was waived because the KNHIS-NSC data set consisted of deidentified secondary data for research purposes. The NHIS is a compulsory healthcare plan for all Korean nationals; eligible citizens are covered either through community- or employee-based plan. The health care utilization DB, one of the main DB run by the Service, was used in the present study. The DB holds a vast amount (over 1.5 trillion cases) of inpatient and outpatient data, including diagnosis, length of inpatient admission, type of treatment, and prescription records.

### Study Cohort

The following criteria were used in search query for extracting benign paroxysmal positional vertigo (BPPV) individuals from the DB; those who 1) had been diagnosed with KCD (Korean Standard Classification of Diseases) Diagnosis Code ‘H811’, and 2) had undergone canalith repositioning procedure (CRP, prescription code ‘MX035’). Atopic dermatitis (AD) individuals were defined as those who 1) had been diagnosed *at least* twice with KCD Diagnosis code ‘L20’ with any two consecutive visits separated by at least 6 months, and 2) had been prescribed topical calcineurin inhibitors (TCI)-tacrolimus (Protopic^®^) 0.03% 10g/0.1% 30g, or pimecrolimus (Elidel^®^) 1% 30g-on the day of diagnosis. Vitiligo individuals were defined as those who 1) had been diagnosed with KCD Diagnosis Code ‘L80’, and 2) had been prescribed topical calcineurin inhibitors, topical corticosteroids-methylprednisolone aceponate 1mg/g (Advantan^®^) 10g, prednicarbate 0.025% (Dermatop^®^) 10mg, *etc.*-or topical calcipotriol 50μg/mL (Daivonex^®^).

### Statistical analysis

A summary of demographic and baseline characteristics was constructed using descriptive analysis: quantitative variables were represented by the mean, maximum, minimum and standard deviation (S.D), while qualitative variables were described by the frequency and proportions (%). Potential associations of BPPV to atopic dermatitis and vitiligo were analysed by χ^2^ tests. Potential disease associations, by sex (male and female), age (≤60, >60), age (≤65, >65), income deciles (0∼4, 5∼7 and 8∼10) and severity grade (normal, moderate and severe), were analysed by χ ^2^ tests. The data analysis was performed by a medical statistician. All statistical analyses were performed using SAS Enterprise Guide 6.1 M1 (SAS Institute Inc., Cary, NC, USA) and IBM SPSS software package for Windows (version 19.0, Chicago, IL). All tests were 2-sided and *p*-values less than 0.05 were ruled statistically significant.

## ACKNOWLEDGEMENTS

This research was supported by a grant of the Clinical Research infrastructure Project through the Korea Health Industry Development Institute (KHIDI) funded by the Ministry of Health & Welfare (H17C2412 to Jong-Yeup Kim). This research was also supported by the Bio & Medical Technology Development Program of the National Research Foundation (NRF) funded by the Korean government (MSIT).

## Author contributions

J.K., D.K., and H.H. designed and conducted the study. H.H. produced the manuscript.

S.K. and J.K. carried out calculations and statistics. All authors read and approved the final manuscript.

H.H. and S.K contributed equally.

D.K. and J.K. contributed equally.

## Data availability

The datasets presented in the current study are available from the corresponding authors upon request.

